# Injury-Specific Muscle Regeneration: A Computational Blueprint for Cellular and Cytokine Drivers

**DOI:** 10.1101/2024.12.16.628612

**Authors:** Megan Haase, Tien Comlekoglu, Alexa Petrucciani, T.J. Sego, Shayn M. Peirce, Silvia S. Blemker

## Abstract

Skeletal muscle regeneration is crucial for maintaining muscle health and mobility, making it a key research focus. Common experimental models of muscle injury used to study this process in vivo include cardiotoxin injury (CTX), freeze-induced (FI) injury, and eccentric contraction (EC) injury. While the response to injury varies across these models, these variations are often overlooked in experimental designs. The cellular dynamics throughout the time course of muscle regeneration differ significantly between these injury types, presenting challenges for both experimental investigation and literature analysis. To enhance understanding of how regeneration responses differ across injury types, we extend our previously validated computational model of skeletal muscle regeneration to simulate muscle fiber remodeling during regeneration following CTX, FI, and EC injuries. We further validate our model against multiple literature-derived metrics of regeneration for each injury type. Analysis of model outcomes reveals that recovery from each injury type is sensitive to unique combinations of cells and cytokines acting at different critical time points, exerting the greatest influence on regeneration at 28 days post-injury. Notably, the cytokines responsible for satellite cell (SSC) recruitment, proliferation, and differentiation required for regeneration vary across injury types, driven by biological redundancy and feedback mechanisms. Specifically, for EC injuries, cross-sectional area (CSA) recovery was predominantly associated with hepatocyte growth factor (HGF) and vascular endothelial growth factor A (VEGF-A) during the early stages of regeneration, with SSC dynamics playing a critical role throughout. FI injuries demonstrated a consistent reliance on HGF across all phases, with additional influences from transforming growth factor beta (TGF-β), tumor necrosis factor-alpha (TNF-α), monocyte chemoattractant protein-1 (MCP-1), and SSC dynamics in later stages. CTX injuries showed early dependence on TGF-β, with significant contributions from SSC, TNF-α, VEGF-A, and fibroblast dynamics over time. Our findings emphasize the importance of considering the molecular and cellular mechanisms relevant to each injury type used in pre-clinical *in vivo* studies, and motivates the development of therapeutic strategies that are designed for specific injury types to optimize recovery outcomes.

## Introduction

Skeletal muscle regeneration occurs as a result of various conditions such as exercise stimuli, contusions, strains, and lacerations^1^. Following initial injury, there is an influx of inflammatory cells and activation of satellite stem cells (SSC) both of which secrete various factors within the microenvironment to signal and regulate cell behaviors^3^. While neutrophils and macrophages clear damaged muscle fibers and apoptotic cells, fibroblasts become activated and deposit collagen to rebuild the extracellular matrix (ECM)^4^. SSCs undergo proliferation, differentiation, and fusion to repair the tissue, and the regeneration cascade begins to shift into a more anti-inflammatory state as macrophages transition into an anti-inflammatory M2 phenotype^5^. Microvascular changes also occur to allow reperfusion of damaged capillaries, further coordinating restoration of the homeostatic microenvironment^6^.

The extent of regeneration varies between injury type; however, the mechanisms for these differences remain unclear. Understanding the distinct mechanisms of muscle injury and regeneration is crucial for interpreting and comparing experimental findings across different studies. However, the use of different muscle injury types and procedures for inducing myonecrosis complicates the interpretation and comparison of results, making it difficult to discern whether outcomes are due to specific treatments and experimental conditions or differences in the initial injury. Common methods for inducing acute muscle injury include eccentric contraction (EC) using maximal lengthening contractions, toxin-induced injury using cardiotoxin (CTX) or barium chloride, and freeze-induced (FI) injury by applying a dry ice-cooled steel probe to the muscle^7–9^. Previously, experimental studies have shown that the trajectory of the regeneration cascade varies with each injury technique^10^. EC injuries typical result from exercise or other forms of high muscular demands, leading to moderate inflammation and faster regeneration than more traumatic injuries^11^. FI results in slower, with prolonged inflammation and all regeneration stages present simultaneously^10^. CTX causes direct muscle fiber necrosis, with synchronous regeneration and a controlled, rapid inflammatory response^12^. Without fully understanding and considering these differences, it becomes challenging to standardize experimental approaches and draw reliable conclusions about muscle regeneration mechanisms across different studies. While some differences between injury types have been found experimentally, many questions still remain: How do the different microenvironments created by each injury type affect the dynamics of SSCs during regeneration? How do different cellular behaviors and signaling factors correlate with muscle fiber recovery for EC, FI, and CTX injuries?

*In silico* computational modeling provides an alternative method for studying specific muscle injury mechanisms comprehensively. Unlike experimental methods, computational approaches allow for the analysis of more frequent timepoints of an individual sample. Computational models have been employed in numerous applications to interrogate the underlying mechanisms responsible for observed tissue dynamics, such as in development^13,14^, disease^15,16^, and wound healing^17,18^. Previously we developed a model of muscle regeneration that simulates the dynamic microenvironment of a cross-section of murine skeletal muscle tissue and robustly calibrated and validated against experimental studies^19^. The goals of this work were to implement different injury types into the existing muscle regeneration model to explore changes in cellular dynamics, microvascular adaptions, and fiber recovery with altered injury conditions.

## Methods

### Agent-Based Model Overview

We utilized the Cellular-Potts modeling framework to develop an agent-based model (ABM) of muscle regeneration in response to skeletal muscle injury as previously described^19^ and summarized in supplemental text 1. Briefly, CompuCell3D (CC3D) version 4.4.1 was used to build a model that spatially represents a two-dimensional female murine skeletal muscle fascicle cross-section. Rules were established based on published studies to govern the behavior of fiber cells, SSCs, fibroblasts, neutrophils, and macrophages, as well as their interactions with the microenvironment. These rules include aspects such as microvasculature remodeling, diffusion, and secretion of regenerative factors like hepatocyte growth factor (HGF), monocyte chemoattractant protein-1 (MCP-1), matrix metalloproteinase-9 (MMP-9), transforming growth factor beta (TGF-β), tumor necrosis factor-alpha (TNF-α), vascular endothelial growth factor A (VEGF-A), interleukin 10 (IL-10), and estradiol (E2). Cell behaviors include recruitment, chemotaxis, phagocytosis, secretion, uptake, activation, proliferation, differentiation, and apoptosis. Unknown parameters were previously calibrated using an iterative parameter density estimation approach described by Haase *et al*^19,20^.

### Inflammatory Cell Agents

Following a prescribed injury, neutrophils and monocytes are recruited via healthy capillaries in response to the altered microenvironmental conditions such as increases in necrosis and MCP-1^21–24^ as well as changes in E2 levels^9,25–27^. Once the neutrophils enter the tissue environment, they chemotax up HGF gradients and phagocytose necrotic tissue, secreting MMP-9, MCP-1, and TNF-α during phagocytosis^22,28–32^. Neutrophils undergo apoptosis after phagocytosis or a pre-defined lifespan elapses^33^. Monocytes chemotax up MCP-1, VEGF-A, TGF-β gradients and can transition into M1 macrophages if TNF-α concentration or a literature defined transition time threshold are met^24,34–38^. Monocytes and M1 macrophages phagocytose necrotic tissue and apoptotic neutrophils and secrete MMP-9, HGF, TGF-β, and IL-10 during phagocytosis^30,38–42^. M1 macrophages can transition to a M2 macrophage phenotype when the M1 has achieved a preset amount of phagocytosis and there is sufficient IL-10 or E2^25,30,42–44^.

### Satellite Stem Cell Agents

SSCs are recruited based on the amount of HGF, MMP-9, and TGF-β within the microenvironment^45–49^. These cells can be activated by local HGF or E2 concentrations and once activated are able to undergo proliferation and differentiation provided that adequate signals are present for the respective behaviors^27,48–52^. To facilitate regeneration, differentiated SSCs fuse to damaged fibers or other differentiated fibers to form new myotubes^2,53–55^. SSCs can return to a quiescent state in the absence of HGF, and can undergo apoptosis when exposed to TGF-beta^48,56^.

### Fibroblast Agents

Fibroblast agents can be activated with sufficient TGF-β at the site of the cell^8,49,57^. Once activated, they migrate towards areas of low collagen and secrete collagen to rebuild the ECM following removal of necrotic tissue by inflammatory cells^49,58,59^. Additionally, they can undergo cell division if near a dividing SSC or transition into myofibroblasts with extended exposure to TGF-β^8,49,60,61^. Following sustained exposure to TNF-α they can undergo apoptosis^49,62^.

### Fiber, Necrotic, Microvascular, and ECM Agents

The morphological agents include muscle fiber cells, necrotic fiber cells, capillaries, lymphatic vessels, and ECM. Collagen densities are prescribed to each ECM agent and varies based on fibroblast/myofibroblast collagen secretion and breakdown of collagen during removal of damaged fibers. The diffusivity of cytokines is altered throughout the simulation based on the ECM collagen density^63^. Following injury, healthy muscle fibers are turned into damaged fiber cells represented by necrosis. Once the necrosis is removed by inflammatory cells it is replaced by low density collagen ECM^22^. Fibroblasts have to secrete collagen at that site to rebuild the ECM prior to placement of a new fiber. Differentiated SSCs can fuse to help rebuild damaged fibers unless the fiber has grown above a certain threshold or the strong cell adhesion between fibers prevents SSC fusion. The contact energies between fibers were set such that the fibers would not merge into unrealistically large fiber masses. Similarly, differentiated SSCs can fuse to each other to form a new fiber if there is sufficient collagen at the site and enough spacing between fibers. Newly formed fibers are assigned a randomly selected maximum size to which the fiber can grow via hypertrophy with additional SSC fusion. Capillaries near areas of necrosis become damaged and must undergo angiogenesis before they can be used as a site for cell recruitment or delivery of signaling factors^64,65^. Lymphatic vessels drain nearby cells and cytokines from the microenvirnonment^66^.

### Muscle histology initialization

The muscle cross-section geometry was created by importing a histology image stained with laminin α2 into a custom MATLAB (The Mathworks Inc., Natick MA) script that masked the histology image to distinguish between the fibers and ECM. The mask was imported into an initialization CC3D script that defined the muscle fibers, ECM, and microvasculature to specific cell types and generated an initialization file that was imported into the ABM as the starting cross-section.

### Incorporation of hypertrophy

Previous iterations of the ABM maintained fiber organization throughout regeneration such that with full regeneration the final fiber configuration matched the starting configuration. Since the introduction of new fibers is unlikely to occur in an identical location and fusion of SSC to existing fibers changes the morphology of the fibers, we have incorporated biologically based rules to include fiber hypertrophy and morphological changes (Fig. 1, animation 1). These rules dictate conserved spacing between fibers and new cell adhesion parameters to maintain ECM between fibers and microvascular structures.

**Figure 1.**
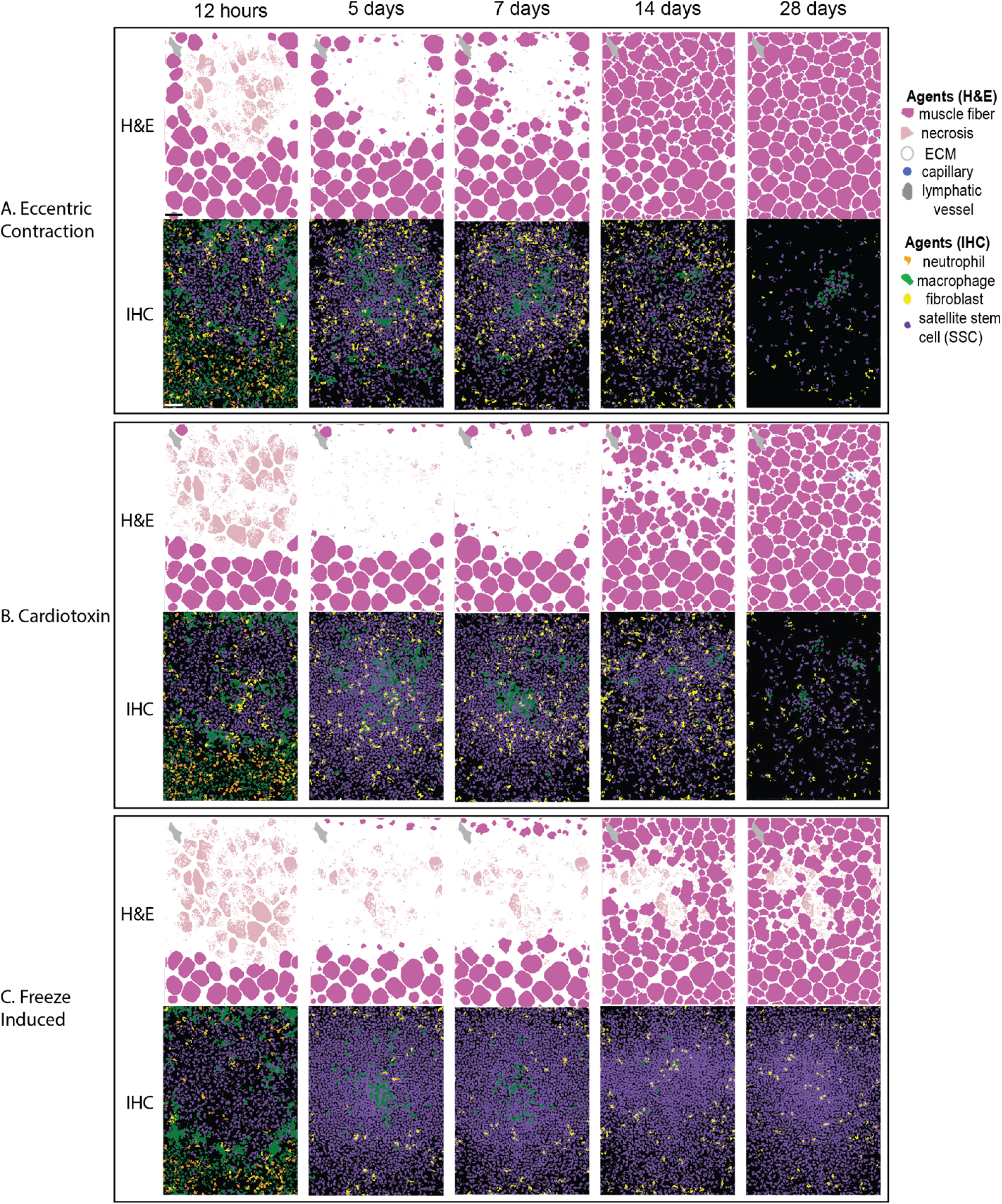
Comparison of fiber cross-section and cellular interaction throughout regeneration simulations for. **(A)** eccentric contraction, **(B)** cardiotoxin, and **(C)** freeze induced injuries highlighting fiber hypertrophy, changes in fiber organization, and spatial cellular changes. Scale bar: 50 µm.

### Injury simulations and recovery outputs

Three different injury types, CTX, FI, and EC injury, were simulated as outlined in Table 1 (Fig. 1, animation 1). Each of these injury conditions were compared to literature experimental data collected from female mice assuming intact female E2 concentration simulation conditions ^7,67,68^.

**Table 1.** Model perturbation input conditions.

| Perturbation | Specific model conditions | References |
| --- | --- | --- |
| Cardiotoxin (CTX) injury | A radius of necrosis was established at the prescribed injury location, leading to complete necrosis of muscle fibers within that area and resulting in approximately 67% cross-sectional damage. Capillaries within the radius of necrosis were also fragmented due to the injury. | 67 |
| Freeze-induced injury (FI) | Clearance of resident SSCs, macrophages, and fibroblasts was achieved by excluding these cells at model initialization, simulating the impaired resident cell population resulting from freeze injury. Damage was localized to a specific section of the cross-section to replicate the contact area of the frozen rod, resulting in approximately 76% cross-sectional damage. Additionally, capillaries within the contact area were fragmented. | 10 |
| Eccentric contraction (EC) injury | A localized radius of necrosis was initialized at a prescribed location resulting in approximately 48% cross-sectional damage, with reduced capillary fragmentation. | 7 |

The literature consistently utilizes the percentage of peak isometric torque as the biomarker for active muscle tissue regeneration^69^. Consequently, in cases where direct cross-sectional area (CSA) measurements were unavailable, model CSA predictions were compared to peak torque measurements from the literature. All conditions were replicated one hundred times to capture the stochastic variability inherent in the model.

### Correlation analysis and random forest regression

To evaluate how regeneration metrics correlated with CSA recovery at the final time point, Spearman’s correlation coefficient was calculated at various time points for each injury type. Additionally, a subset of these metrics was used to train a random forest regression model. The random forest model was iterated three hundred times to ensure robustness, and the average R^2^ and permutation importance were calculated for each injury type. After averaging permutation importance scores across iterations, we selected the top 40% of features for the final random forest model.

### Statistics

Comparisons between groups were performed using a multiway ANOVA in MATLAB with significant p-values ≤.05 unless otherwise noted. The results were analyzed with 95% confidence intervals.

## Results

### ABM outputs agree with experimental data across various injury conditions and outputs

The model’s simulations of muscle regeneration for each injury type generally align with the literature. Model-predicted recovery in regenerating minimum fiber diameter following CTX injury aligned with experimental data (Fig. 2A). Freeze injury model data were consistent with experimental trends for CSA recovery, with the 95% confidence interval within the standard deviation for all time points (Fig. 2B). For EC injury, model simulations fell within literature CSA recovery ranges at all reported time points, predicting muscle hypertrophy above 100% of the original fiber CSA beyond 14 days post-injury (Fig. 2C). However, both FI and EC had a time point where the mean model prediction was outside of the experimental range.

**Figure 2.**
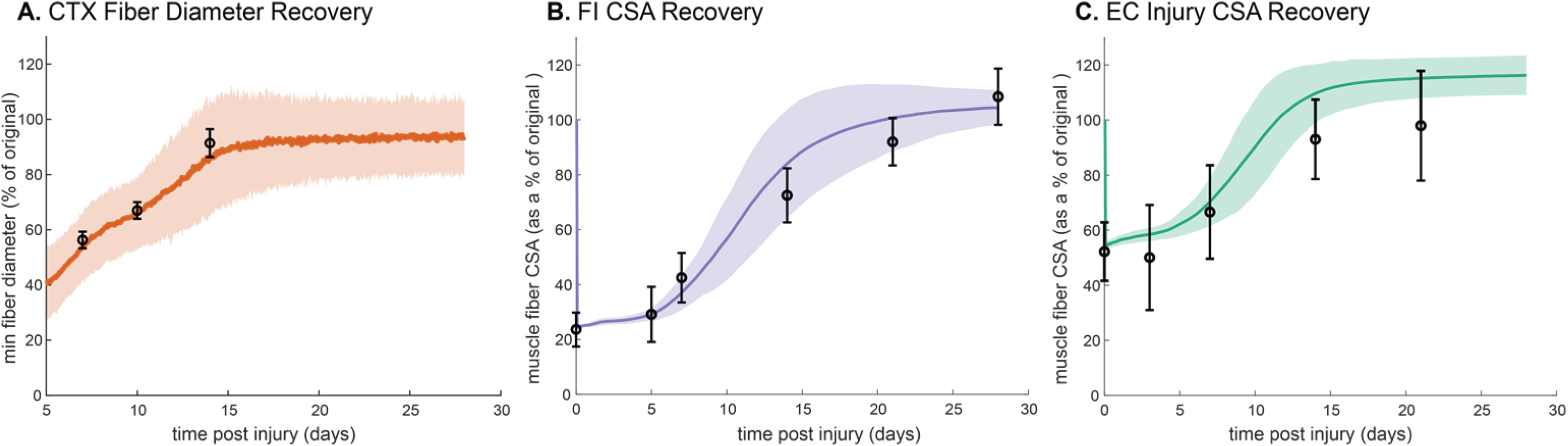
ABM injury condition validation. Model fiber recovery metrics were compared to experimental data for cardiotoxin. (A)^75^, freeze injury (B)^9^, and eccentric contraction injury (C)^76^. Error bars represent experimental standard deviation, and model 95% confidence interval is indicated by the shaded region.

Model predicted differences in SSC count, vascularization, fiber diameter, necrosis, MCP-1, and macrophages between injury types were compared at timepoints reported experimentally.

Predicted statistical changes between conditions were assessed against literature results, and model-predicted changes between injury types were found to be consistent with experimental data (Table 2, Table 3). CSA recovery occurred fastest and with the most hypertrophy in the EC simulation, followed by the CTX injury (Fig. 3A), consistent with prior experimental findings^10,11^. The FI condition resulted in delays in CSA recovery but reached similar myofiber diameters by 28 days post injury as the CTX injury (Fig. 3B), also aligning with previously reported results^10^. Model capillary counts were significantly different between each injury type (Fig. 3J). At 28 days post injury the EC injury resulted in the highest number of capillaries, and the least in simulated FI, similar to experimental results which have shown FI results in highest rates of vascular destruction^10^. M1 peaks were similar between CTX and FI, but CTX showed a later peak compared to FI (Fig. 3H) consistent with literature results showing an early induction of genes mediating inflammatory infiltration in FI^11^.

**Figure 3.**
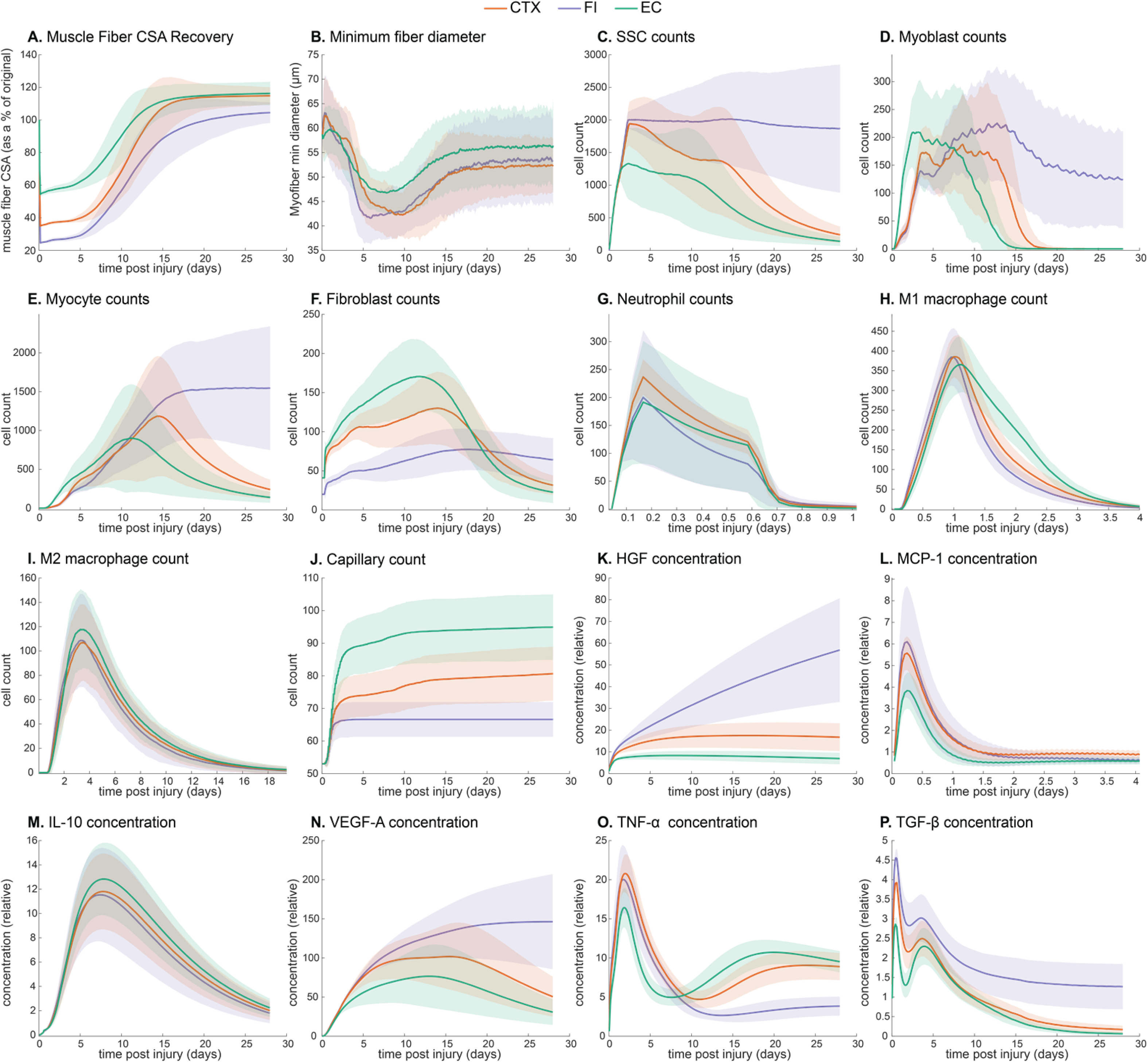
Comparison of regeneration metric outputs for freeze injury, cardiotoxin, and eccentric contraction.

**Table 2.** Model validation of additional metrics comparing outputs between FI and CTX injury conditions^10^ (NS, not significant change)

|  | FI |  | CTX |  |
| --- | --- | --- | --- | --- |
|  | Model | Experiment<sup>10</sup> | Model | Experiment<sup>10</sup> |
| SSC (2 days) | ↑ | ↑ | ↓ | ↓ |
| Capillaries (4 days) | ↓ | ↓ | ↑ | ↑ |
| Fiber diameter (28 days) | NS | NS | NS | NS |
| Necrosis (4 days) | ↑ | ↑ | ↓ | ↓ |

**Table 3.** Model validation of additional metrics comparing outputs between FI and EC injury conditions^11^.

|  | **FI** |  | **EC** |  |
| --- | --- | --- | --- | --- |
|  | Model | Experiment<sup>11</sup> | Model | Experiment<sup>11</sup> |
| M1 (6 hrs) | ↑ | ↑ | ↓ | ↓ |
| CSA (4 days) | ↓ | ↓ | ↑ | ↑ |
| MCP-1 (6 hrs) | ↑ | ↑ | ↓ | ↓ |

**Table 4.** Random forest model features used for each injury type highlighting importance of metrics for random forest regression model prediction.

| CTX |  |  | FI |  |  | EC |  |  |
| --- | --- | --- | --- | --- | --- | --- | --- | --- |
| Metric | Day | Feature importance | Metric | Day | Feature importance | Metric | Day | Feature importance |
| Myocytes | 14 | 0.35039 | Myocytes | 21 | 0.52203 | SSCs | 2 | 0.31635 |
| SSCs | 14 | 0.34697 | MCP-1 | 21 | 0.45878 | SSCs | 21 | 0.31374 |
| TNF-α | 21 | 0.32893 | TGF-β | 21 | 0.34664 | VEGF-A | 5 | 0.29497 |
| TNF-α | 14 | 0.3226 | HGF | 21 | 0.3208 | VEGF-A | 7 | 0.28383 |
| VEGF-A | 21 | 0.29206 | TNF-α | 21 | 0.29459 | Myocytes | 14 | 0.28192 |
| VEGF-A | 14 | 0.26755 | HGF | 14 | 0.28784 | MMP | 21 | 0.26216 |

### Model predicts injury-specific cellular changes and microenvironmental signaling profiles

The total SSC count showed similar peaks and timings across all injury conditions, but the FI condition resulted in prolonged elevation of SSCs with more variability in cell counts compared to the other injury conditions (Fig. 3C). There was a significant difference among all injury simulations in SSC differentiation into myoblasts and myocytes. The EC simulation exhibited the earliest and highest peak in myoblasts and myocytes, followed by an earlier decrease compared to FI and CTX injuries (Fig. 3D, E). Similar but lower trends were observed for myoblasts and myocytes with CTX injury, whereas the FI simulation showed a greater delay and remained elevated at later stages of regeneration. Fibroblast cell counts and CSA recovery were significantly different between each of the injury simulations (Supplemental table 1). Fibroblast counts during the intermediate phase of regeneration (5-15 days post injury) were greatest in EC injury and least in freeze injury (Fig. 3F). Neutrophil peaks were highest with CTX compared to other injuries (Fig. 3G). M2 peaks were highest with EC, followed by CTX and then FI (Fig. 3I).

Simulated MCP-1, HGF, and TGF-β cytokine mean field values were greatest in the FI and least in the EC injury mechanism (Fig. 3K, L, P). IL-10 and VEGF-A mean field values were similar throughout regeneration for the various injury mechanisms, however, mean VEGF-A field values plateaued towards the end of regeneration (20 days post injury) with the FI injury whereas they decreased after 15 days post injury in the CTX and EC injury mechanisms (Fig. 3M, N). TNF-α field values were lowest in EC injury before 9 days post injury, and greatest after 10 days post injury (Fig. 3O). Mean TNF-α values for the FI injury were similar to those of the CTX injury before 10 days post injury and remained lower than the other two injury mechanisms after 10 days post injury (Fig. 3O).

### Correlations between regeneration metrics and CSA recovery exhibit temporal and injury dependence

Spearman’s correlation between regeneration metrics and CSA recovery at day 28 illustrated that distinct regeneration patterns in FI, CTX, and EC injuries are primarily driven by differing activities of fibroblasts, SSCs, macrophages, and alterations in concentrations of HGF, TGF-β, and TNF-α (Fig. 4A-C). In FI, fibroblasts consistently showed positive correlations, peaking at day 14 (Fig. 4A), whereas CTX had variable correlations, shifting from negative to positive at days 7 and 14, then back to negative by day 21 (Fig. 4B). EC exhibited positive fibroblast correlations between days 5 and 14, but negative on day 2 and 21 (Fig. 4C). SSCs in FI switched from weak negative correlations to positive at days 7 and 14, then back to negative, while in CTX, SSCs maintained strong positive correlations from days 5 to 21. EC displayed consistent positive SSC correlations. M1 and M2 macrophages were positively correlated in FI early on, but negatively correlated in CTX throughout. In EC, macrophages showed negative correlations at early timepoints but positive correlations at later timepoints. HGF was negatively correlated in FI but positive in both CTX and EC. TGF-β had a consistent negative correlation in FI but varied in CTX and EC. TNF-α, VEGF-A, IL-10, and MCP-1 showed distinct temporal correlation patterns across the injuries. These results highlight that fibroblast and macrophage activities, along with the signaling molecules HGF, TGF-β, and TNF-α, are pivotal in distinguishing the regenerative mechanisms between the injury types.

**Figure 4.**
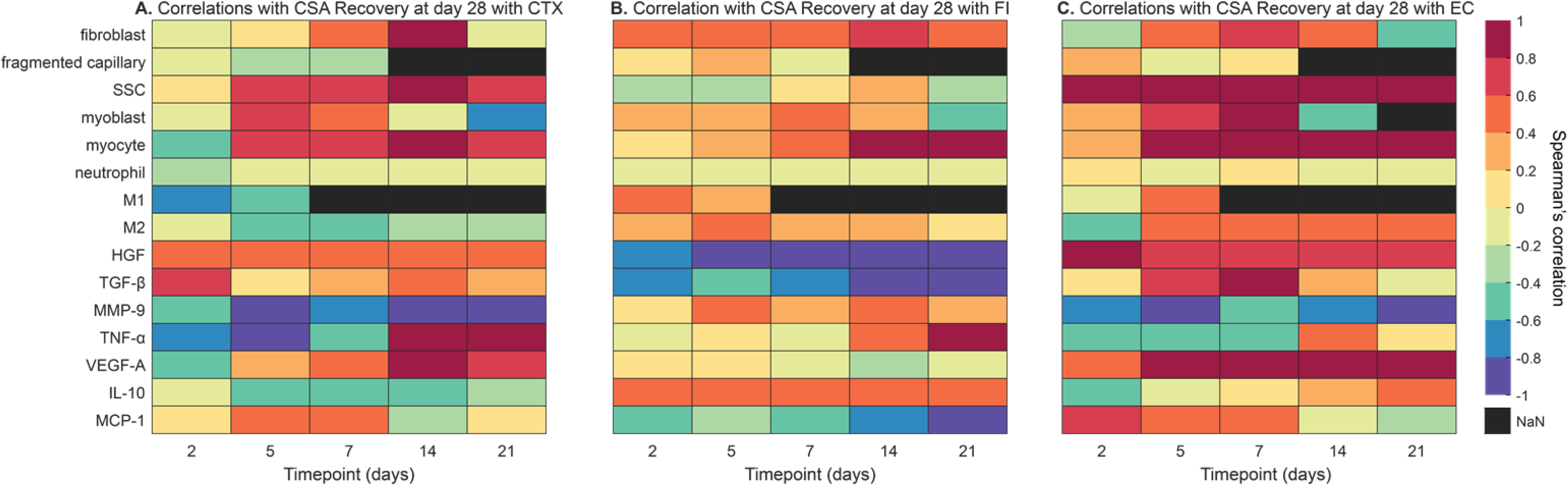
Heatmap of correlations between various regeneration metrics and CSA recovery at 28 days post cardiotoxin injury (A), freeze injury (B), and eccentric contraction (EC).

### Cytokines regulating SSC dynamics necessary for regeneration vary by injury type

Correlation heatmaps revealed that the cellular factors influencing injury outcomes after 28 days varied across injury types (Figure 4). For EC injuries, CSA recovery was prominently linked to HGF and VEGF-A during early regeneration stages, with significant contributions from SSC dynamics throughout. In contrast, for FI injuries, CSA recovery was most correlated with HGF across all regeneration phases, along with additional influences from TGF-β, TNF-α, MCP-1, and SSC dynamics in later stages. For CTX injuries, CSA recovery was positively correlated with TGF-β during early regeneration, with SSC dynamics becoming more significant in later phases, accompanied by TNF-α, VEGF-A, and fibroblast dynamics. These findings indicate that different injury types initiate unique combinations of cytokines and growth factors to recruit SSC behaviors essential for regeneration.

A random forest model was developed to identify the most significant regeneration metrics predicting CSA recovery across different injury types. Initially, the model incorporated all available metrics, but to enhance accuracy and prevent overfitting, features with low permutation importance were excluded, retaining only the top 40% of the most impactful predictors. Employing these identified features yielded R^2^ values of 0.611 for EC, 0.761 for CTX, and 0.657 for FI. The permutation importance’s for each injury condition, highlighting which metrics contributed heavily to prediction accuracy, are included in Table 3. For FI, the most important feature was myocytes at day 21, with an importance score of 0.522, followed by MCP-1 concentration at day 21 (importance: 0.459) and TGF-β concentration at day 21 (importance: 0.347). For CTX, the most important features were myocytes at day 14 (importance: 0.350), SSCs at day 14 (importance: 0.347), and TNF-α concentration at day 21 (importance: 0.329). For EC, SSCs at day 2 was the most important feature (importance: 0.316), followed by SSCs at day 21 (importance: 0.314), and VEGF-A concentration at day 5 (importance: 0.295). These results underscore that cytokines influencing SSC dynamics vary across different injury types, highlighting the multiple pathways and signals involved in regeneration for each type of injury.

## Discussion

Comparative studies have highlighted distinct differences among commonly used injury mechanisms for studying muscle regeneration after acute injury. However, understanding how recovery is mediated across different injury mechanisms remains challenging. In this work, we provide comprehensive insights into how the distinct microenvironments created by different types of injuries influence SSC dynamics during muscle regeneration and correlate with muscle fiber recovery. Our model simulations align closely with experimental data across various injury conditions, revealing that each injury type elicits unique patterns of cellular behaviors and microenvironmental signaling. Importantly, our findings underscore that muscle can regenerate through multiple mechanisms, with different signaling factors becoming more prevalent depending on the injury type. These variations arise from the differing ways in which SSC dynamics are triggered and sustained to elicit regeneration. For example, FI injury prolongs SSC elevation with greater variability in cell counts, while EC injury shows early and robust peaks in myoblasts and myocytes, followed by an earlier decline compared to CTX and FI injuries. These distinct SSC dynamics, along with the temporal changes and concentrations of MCP-1, HGF, TGF-β, TNF-α, and VEGF-A, significantly contribute to the varied rates and extents of muscle fiber recovery observed across different injury types. The redundancy and feedback mechanisms inherent in these biological processes further emphasize the complexity of muscle regeneration and the need to tailor therapeutic strategies to specific injury contexts. *Impact of Injury Condition Variations on Regeneration Outcomes*

Our computational model aligns with previously published experimental findings that highlighted significantly different pathophysiologic mechanisms across different injury types^10^. While CTX, FI, and EC are commonly used in muscle regeneration research, the rationale for selecting a particular injury type is often inadequately justified. This work underscores two critical findings: first, it elucidates distinct groups of cellular and biochemical mechanisms underlying regeneration from specific injuries, emphasizing the need to tailor experiments and therapeutic strategies accordingly. Second, it highlights the utility of computational models in probing these injury-specific mechanisms through hypothesis generation and testing. Furthermore, the impact of different injury types is often overlooked when interpreting and comparing results related to regeneration metrics or treatments. This oversight is a significant confounding variable that contributes to contradictory findings regarding treatment outcomes^70,71^. Given the varying microenvironmental conditions, cellular behaviors, and regeneration trajectories associated with each injury condition, both computational models and experimental designs must carefully consider the selection of the injury type based on the specific research being addressed.

### ABM elucidates temporal injury dependent cellular mechanisms

Comparative work analyzing cell counts and gene expression profiles at multiple time points post injury revealed significant differences among the injury types over the time course of regeneration^11^. Our work confirms these differences and identifies unique cellular, cytokine, and microvasculature changes that influence regeneration for each injury type. We note particularly prolonged elevation of HGF in later stages of regeneration in FI as compared with CTX and EC injuries, suggesting greater inflammatory signaling as CSA recovers in this injury mechanism. We observe delayed CSA recovery in FI compared with the other injury types despite the greatest SSC counts and prolonged HGF levels unique to this condition, indicating a potential oversaturation of SSCs that could inhibit regeneration in this injury mechanism. There may be an intermediate level of SSCs and HGF that is most beneficial for regeneration that is reached for CTX and EC injuries but not for FI. Additionally, despite prolonged VEGF-A levels, FI cases failed to reach the elevated capillary counts observed in CTX and EC injuries.

Regression analysis further demonstrated differences in the regeneration time course of the injury mechanisms. By identifying correlations between regeneration metrics and CSA recovery at day 28, along with feature importance’s from our predictive models, we were able to gain a deeper understanding of the key biological processes involved in muscle healing for each injury type. For EC injuries, CSA recovery was predominantly associated with HGF and VEGF-A during the early stages of regeneration. The significance of these signaling factors highlights their roles in promoting angiogenesis and tissue repair soon after injury, consistent with previous findings^72,73^. Additionally, SSC dynamics were found to play a critical role throughout the recovery process and were associated with CSA recovery outcomes 28 days post injury. These findings suggest that early intervention strategies targeting these factors could be beneficial in enhancing recovery outcomes for EC injuries^74^. For FI injuries, analysis revealed consistently strong negative correlations between HGF and CSA recovery at 28 days post injury. Additionally, TGF-β, TNF-α, MCP-1, and SSC dynamics were influential in the later stages of recovery. These results emphasize the multifaceted nature of muscle repair in FI injuries, suggesting that therapeutic approaches should consider multiple factors over an extended period. For CTX injuries, TGF-β was particularly influential during the early stages of regeneration, while SSC dynamics became more significant in the later phases. Additionally, TNF-α, VEGF-A, SSC, and fibroblast dynamics were important throughout the recovery process. These findings highlight the critical window of early intervention and the ongoing need to support regenerative processes throughout muscle healing. Correlation and permutation importance analysis underscored the heterogeneity in the factors influencing CSA recovery across different injury conditions. Each injury type exhibited distinct sets of critical features, emphasizing the need for tailored therapeutic strategies. The feature importance highlighted distinct features as predictors for each injury type, emphasizing the heterogeneity in the factors influencing CSA recovery across different injury conditions. The identification of specific features at particular time points provides valuable insights into the temporal dynamics of muscle regeneration and offers potential targets for enhancing recovery outcomes.

### Leveraging specific injury types to study mechanisms of treatment interventions

The computational model provides novel insight into the differences between mechanisms for each injury type and which regeneration metrics are most crucial for each injury. During the experimental design process, our model could be leveraged to identify which injury type is most suitable to explore specific mechanisms and interactions during regeneration. Furthermore, our model indicates that injury type is likely an important factor when selecting and formulating a treatment strategy, since both the significance and timing of candidate therapeutic mechanisms vary by injury type. For example, if evaluating the delivery of differentiated SSCs for improving CSA recovery, FI would likely not have beneficial outcomes because the microenvironment is already over-saturated with SSCs. Beyond assisting in injury type selection, the model could also be used as a tool to inform optimal treatment delivery timing based on the correlations between specific mechanisms and CSA recovery.

### Limitations and future directions

It is important to recognize some key limitations of the model. First, the use of only one region of muscle tissue limits the recruitment of inflammatory cells from nearby or injury-adjacent tissue. Scaling up the model to include multiple compartments or sections from the muscle tissue would be beneficial for examining higher-grade injuries and their impact on surrounding tissue. It is also important to note that the model is based on female histology and calibrated using female data. Future work could extend the model to incorporate sex differences and how they might impact injury response. Additionally, we utilized only one initial histology configuration, although the model can be adapted to incorporate various histology images^19^. This flexibility could be applied in future applications to explore how initial fiber morphology influences regeneration outcomes. Another limitation is the assumption of a consistent injury amount for each injury type, despite recognizing the variability in injury extent depending on the protocol used. Future research could utilize this model to investigate how variations in injury delivery impact results and how different methods of inducing the same injury might lead to divergent outcomes.

Additionally, future studies should explore the impact of injury location, considering that different tissues may be more sensitive to specific injuries. The model could also serve as a valuable tool for determining which injury type is most appropriate for specific experiments and for comparing studies that use different injury types. This approach would be particularly useful for comparing experiments that test treatments using different injury models, helping to account for the confounding effects of injury variability.

This study underscores the importance of accounting for injury condition differences when investigating muscle regeneration and highlights the unique cellular and molecular mechanisms that underlie recovery from different injury types. By using an advanced computational model, we replicated and analyzed the regeneration processes following three common skeletal muscle injuries used in pre-clinical research, demonstrating the model’s ability to predict cellular dynamics consistent with experimental findings. Our results reveal that muscle regeneration can occur through multiple mechanisms, with different signaling factors becoming more prevalent depending on the injury type. These variations are driven by the differing methods through which SSC dynamics are triggered and sustained, influenced by the unique microenvironments and biological redundancy inherent in muscle regeneration. The model emphasizes the critical influence of distinct factors at specific stages of regeneration on CSA recovery, suggesting that targeted therapeutic strategies should be tailored to the specific injury type to optimize muscle healing. Furthermore, this model can serve as a valuable tool to bridge the gap between various mouse models and human muscle injury, enhancing the translational potential of pre-clinical findings. This work provides valuable insights into the temporal dynamics of muscle regeneration and offers a robust framework for future research to investigate the impact of injury variations. Ultimately, it aids in the development of more effective treatment interventions by considering the complexity and variability of muscle regeneration processes across different injury contexts.

## Acknowledgements

The authors acknowledge NIH Grants #R21AR080415 and #U24EB028887, Wu-Tsai Foundation Agility Project Funding, Achievement Rewards for College Scientists, and NSF GRFP Grant #1842490 for financially supporting this research.

## Additional Resources

https://simtk.org/projects/musinjuriesabm

Large animation, see separate file

**Animation 1.**
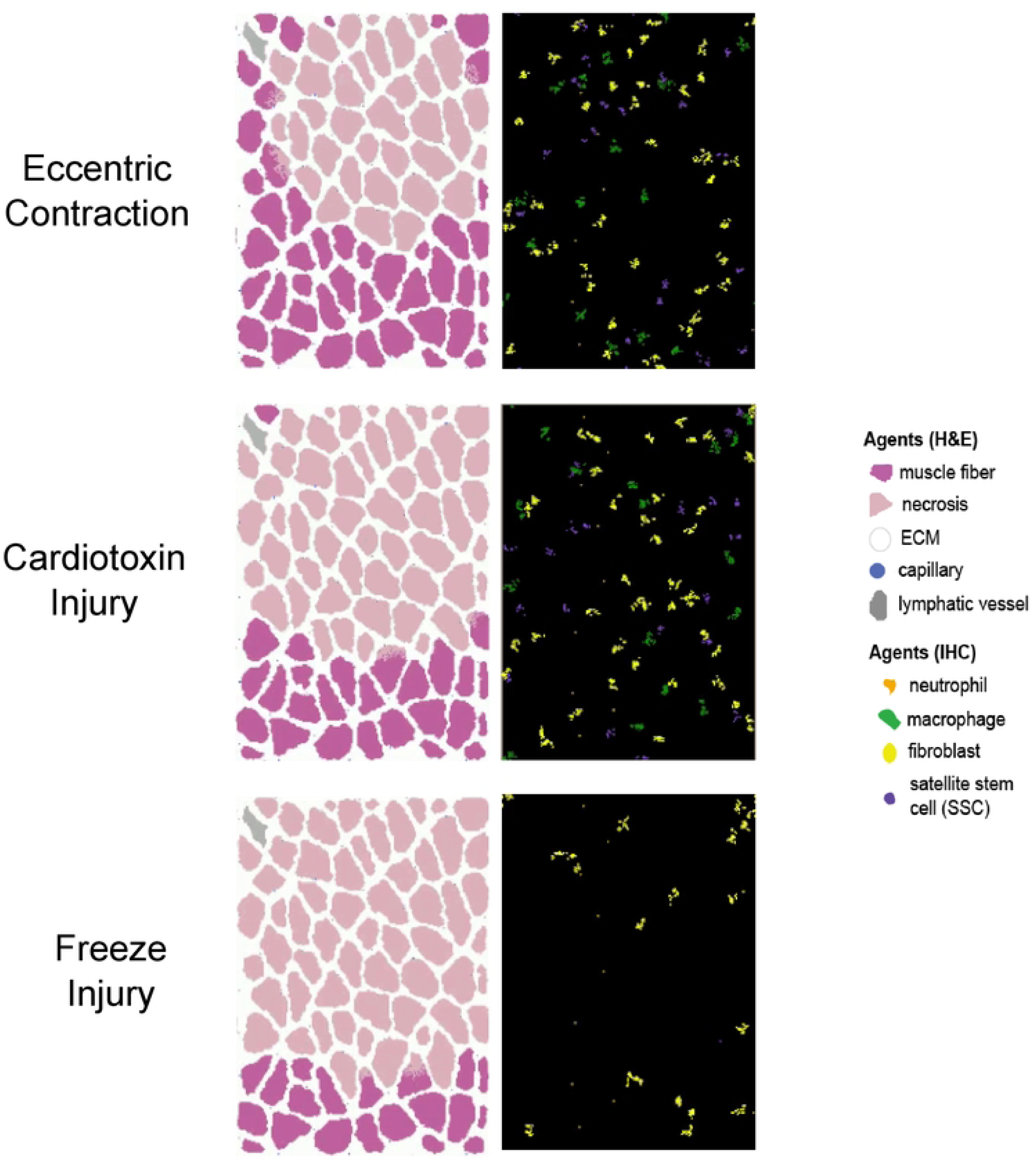
Animation of fiber cross-section and cellular interaction throughout regeneration simulation highlighting fiber hypertrophy, changes in fiber organization, and spatial cellular changes.

**Supplemental Table 1.** P-values from injury comparison ANOVAs.

| Metric | D2 |  |  | D5 |  |  | D7 |  |  | D14 |  |  | D28 |  |  |
| --- | --- | --- | --- | --- | --- | --- | --- | --- | --- | --- | --- | --- | --- | --- | --- |
|  | FI vs CTX | FI vs EC | CTX vs EC | FI vs CTX | FI vs EC | CTX vs EC | FI vs CTX | FI vs EC | CTX vs EC | FI vs CTX | FI vs EC | CTX vs EC | FI vs CTX | FI vs EC | CTX vs EC |
| CSA | 0 | 0 | 0 | 0 | 0 | 0 | 0 | 0 | 0 | 0 | 0 | 1.359 E-07 | 0 | 0 | 0.005 |
| Min fiber diameter | 0.001 | 0.002 | 0.833 | 0 | 0 | 0.001 | 3.849 E-07 | 0 | 5.590 E-18 | 0.915 | 5.870 E-14 | 2.050 E-15 | 0.136 | 5.128 E-07 | 2.617 E-12 |
| SSC | 2.357 E-08 | 0 | 0 | 1.188 E-06 | 0 | 0 | 9.072 E-23 | 0 | 0 | 0 | 0 | 0 | 0 | 0 | 0.035 |
| Myoblast | 0.380 | 0 | 0 | 3.765 E-08 | 0 | 6.626 E-16 | 0.018 | 0.929 | 0.006 | 0 | 0 | 0 | 0 | 0 | 1 |
| Myocyte | 0.1295 | 0 | 0 | 1.064 E-23 | 0 | 5.863 E-17 | 0.0003 | 0 | 1.174 E-11 | 0.045 | 0 | 2.241 E-22 | 0 | 0 | 0.007 |
| Fibroblast | 0 | 0 | 0 | 0 | 0 | 0 | 0 | 0 | 0 | 0 | 0 | 0 | 0 | 0 | 1.821 E-10 |
| Neutrophil | 9.192 E-07 | 3.384 E-23 | 7.679 E-06 | 1.222 E-06 | 4.145 E-14 | 0.0248 | 1.143 E-10 | 5.549 E-16 | 0.257 | 1.435 E-13 | 1.597 E-17 | 0.545 | 7.538 E-15 | 3.984 E-19 | 0.535 |
| M1 | 0 | 0 | 0 | 0.125 | 0.159 | 0.0005 | N/A | N/A | N/A | N/A | N/A | N/A | N/A | N/A | N/A |
| M2 | 0.001 | 0.0003 | 0.920 | 0.029 | 1.434 E-13 | 2.045 E-06 | 8.420 E-05 | 1.057 E-15 | 0.0003 | 0.014 | 3.138 E-08 | 0.010 | N/A | N/A | N/A |
| Capillaries | 0 | 0 | 0 | 0 | 0 | 0 | 0 | 0 | 0 | 0 | 0 | 0 | 0 | 0 | 0 |
| MCP-1 | 8.759 E-10 | 0 | 0 | 0 | 0.0001 | 0 | 0 | 0 | 0 | 3.545 E-24 | 0.009 | 6.109 E-12 | 0 | 0 | 0.126 |
| HGF | 0 | 0 | 0 | 0 | 0 | 0 | 0 | 0 | 0 | 0 | 0 | 0 | 0 | 0 | 4.003 E-22 |
| TGF- $\beta$ | 0 | 0 | 0 | 0 | 0 | 0.007 | 0 | 0 | 0.101 | 0 | 0 | 1.395 E-09 | 0 | 0 | 3.351 E-05 |
| TNF-α | 0.0001 | 0 | 0 | 0 | 0 | 0 | 1.405<br>E-22 | 0 | 0 | 0 | 0 | 0 | 0 | 0 | 3.543<br>E-09 |
| IL-10 | 4.575<br>E-06 | 3.559<br>E-19 | 9.874<br>E-05 | 0.594 | 0.008 | 0.0002 | 0.712 | 1.380<br>E-06 | 6.189<br>E-05 | 0.010 | 5.739<br>E-13 | 2.996<br>E-05 | 0.0004 | 1.235<br>E-14 | 0.0002 |
| VEGF-A | 0.002 | 0 | 0 | 0.001 | 0 | 0 | 3.945<br>E-08 | 0 | 0 | 0 | 0 | 3.131<br>E-22 | 0 | 0 | 4.341<br>E-12 |

## Supplemental Text 1. Cellular-Potts

### Cellular-Potts model implementation

In the CPM, individual cells are represented as a collection of pixels on a square lattice. These computational representations of cells have properties of predefined volume, contact energy with surrounding cells in their environment, and affinity or aversion to diffusing species that drive chemotactic behaviors^77^. These properties are defined mathematically in the Effective Energy function *H* which is evaluated in every computational timestep. Equation (1) represents the effective energy function that imposes the physical properties of all cells in the agent-based CPM.

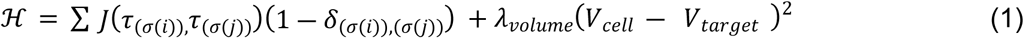

The first term describes contact energy of each cell for neighboring cells in its environment governed by *J* the contact coefficient, where *i,j* denote neighboring lattice sites, τ denotes cell types, and σ denotes individual cells in the simulation. The delta function localizes contact energy contributions to cell-cell interfaces. The second term represents a volume constraint scaled by λ_*volume*_ on each cell. Motile cells in the simulation that exhibit chemotactic behavior towards a target have additional terms to drive their movement described in (2).

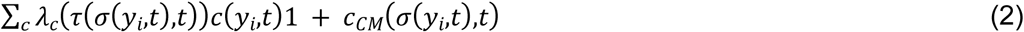

Which models logarithmic chemotaxis by cell type and chemokine field concentration influenced by chemotaxis parameter λ_*c*_, chemical field concentration *c*, and cell body center-of-mass measurement *c*_*CM*_from which chemotaxis behaviors are calculated. Diffusion and secretion behavior of chemical species is governed by the general diffusion equation described in (3).

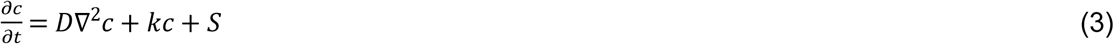

Where *k* defines the decay constant of diffusive species concentration *c*, for which *D* is the diffusion constant. The *S* term is an additional term to represent secretion of diffusive species.

To apply these relations to recreate cell movement, the CPM randomly selects a pair of neighboring pixels and evaluates whether one pixel may copy itself to the location of the other, called a “copy attempt.” The probability of acceptance or rejection of that pixel copy attempt is governed described by the Boltzmann acceptance function described in (4).

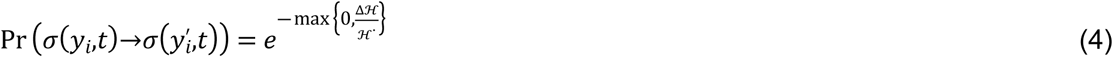

This function describes the probability of the copy attempt 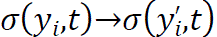 where pixel 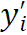 copies to pixel *y*_*i*_in the CPM algorithm^77^. The value ℋ^∗^ represents our *temperature* parameter in the CPM, or the parameter that affects stochasticity of lattice site copy attempts. Δℋ is the change in effective energy due to a lattice site copy attempt. The Cellular-Potts simulation discussed here was implemented using CompuCell3D (CC3D), an open-source Python-based CPM platform^77^.

### Cellular-Potts model initialization

The muscle cross-section geometry was initialized by importing a histology image of (murine gastrocnemius stained with laminin α2) masked to distinguish between muscle fiber and ECM. The mask was imported into an initialization CC3D script that defined the muscle fibers, ECM, and microvasculature to specific Cellular-Potts lattice types and generated a Potts Initialization File (PIF) that was imported into the ABM as the starting cross-section. This initial configuration resulted in a square lattice of dimensions 321×417 pixels. The model consisted of 2 layers in the z axis: a tissue environment layer consisting of muscle, ECM, capillaries, and lymphatic channels, and an immune cell layer where immune cell agents migrated and interacted in the “2.5D” model. CPM interaction neighbor order was 4 for typical adhesion energy terms, and 1 for all other CPM Hamiltonian terms.

